# Transcriptomic Characterization of Terminal Complement Complex-bound Cells in Human Choroid Using Single-Cell RNA Sequencing

**DOI:** 10.64898/2026.09.05.749603

**Authors:** Renato D Jensen, Nathaniel K Mullin, Kelly Mulfaul, Jack EB Miller, Emma Navratil, Andrew P Voigt, Todd Scheetz, Luke A Wiley, Edwin M Stone, Budd A Tucker, Robert F Mullins

## Abstract

Age-related macular degeneration (AMD) is among the leading causes of blindness worldwide. Early AMD is characterized by dysfunction in the choroid, including early dropout of endothelial cells and increased deposition of the complement cascade’s membrane attack complex (MAC) in the choriocapillaris. In this study, we used a single-cell RNA sequencing-based approach with barcoded antibodies to measure abundance of the MAC and other surface proteins at single cell resolution on RPE/choroid samples from four aged human donor eyes. We included antibodies to detect the MAC, CD34, CD45, and complement regulators CD55 and CD59, in addition to control antibodies. Our analysis of these data revealed cell clusters with expected gene expression profiles and antibody-based detection of CD34 and CD45 congruent with transcriptome-based cell identity. We also detected surface complement regulators CD55 and CD59 across a wide variety of cell types. Across endothelial cells, surface CD55 and CD59 appeared more abundant on venous clusters, and abundance of each was correlated with expression of a third complement regulator, clusterin (*CLU*). The MAC was detected on a variety of cell types, but was most abundant on the surface of cells in the macrophage family, smooth muscle cells, and pericytes. We confirmed these findings by identifying MAC deposition on choriocapillaris pericytes using immunohistochemistry for MAC, endothelial, and pericyte markers. Ultimately, these data showcase a valuable new approach to analyze gene expression and surface complement in human donor eyes, and provide novel insight into patterns of MAC deposition and complement protection in the aging human choroid.

## Introduction

Age-related macular degeneration (AMD) is among the leading causes of blindness worldwide [1,2,3]. It is a disease of the retina and choroid that affects high-acuity, central vision in the growing aging population [4]. There are no good treatments for the early stages of AMD, and few available for its potentially blinding later stages [5]. Despite high disease prevalence and significant impact on vision and quality of life, the inciting pathophysiology remains unclear. Although age is by far the most impactful risk factor for developing AMD, a common polymorphism (Y402H) in *CFH*, a gene encoding a regulator of the innate immune system’s complement cascade (factor H), increases disease risk up to 7-fold in homozygotes, and at least one copy of this polymorphism is seen in almost 50% of patients with AMD [6,7]. Other mutations in genes encoding complement proteins and their regulators are also associated with modified risk for developing AMD [8,9]. Complement dysregulation has been noted in the choroid and RPE of patients with early AMD. For example, the C5b-9 membrane attack complex (MAC) shows increased abundance in the choriocapillaris in eyes with AMD and in eyes with high risk CFH genotypes [10, 11]. Furthermore, there is histopathological evidence that endothelial cells in the choriocapillaris die early in the course of disease [12]. Accordingly, understanding the impact of the complement cascade on the choroidal endothelium with age and in AMD may offer new translational insights.

Some complement regulators implicated in modulating AMD risk, like factor H, are secreted and circulate or bind to extracellular matrix [13]. However, cells also produce surface-bound proteins that inhibit activation of the complement cascade at the cell membrane. One such protein is CD55 (decay-accelerating factor, or DAF), which is present on the cell surface and inhibits complement activation by accelerating decay of C3 and C5 convertases [14,15].

Another is CD59, a terminal complement inhibitor, which prevents assembly of the membrane attack complex (terminal complement complex, C5b-9, or MAC) on the cell membrane. The stepwise nature of choroidal endothelial cell loss suggests that certain cells are more susceptible to the inciting pathophysiology of AMD than others, but the molecular basis for this susceptibility remains unclear. In the context of the numerous complement-based risk factors for AMD, we sought to evaluate the relationship between abundance of surface-bound complement regulators and gene expression in choroidal microvascular cells.

Notably, the complement cascade’s MAC has been shown to accumulate in the choriocapillaris with age, and is more abundant in the choriocapillaris of AMD patients [10]. However, quantifying MAC abundance across different cell types in the eye has remained difficult. Technologies like single-cell RNA sequencing (scRNA-seq) have allowed for powerful gene expression-based comparisons across numerous cell types in parallel [16]. However, at least some components of the MAC deposited in the choroid are primarily produced in the liver and cannot be detected in the gene expression of its components by ocular cells. Accordingly, the application of this powerful tool to assay many cells in parallel for surface MAC necessitates the use of a multiomic approach combining gene expression and cell surface protein.

Recently, single-cell RNA sequencing has been paired with oligonucleotide barcoding of antibodies, allowing for correlation of cell surface proteins with global gene expression [17]. The aim of this study was to characterize membrane-bound MAC and its regulators across the various cell types of the human choroid in an unbiased manner by combining single-cell RNA sequencing with cell surface protein detection. We sought to establish use of this multiomic approach in human ocular tissue to evaluate the patterns of complement deposition and susceptibility in the aging choroid, with potential implications for the pathophysiology of early AMD.

## 1. Materials and Methods

### Human Donor Eyes

Fifteen human donor eyes were utilized for the various experiments in this study. Donations were acquired through the Iowa Lions Eye Bank with complete consent by donors’ next of kin and in accordance with the Declaration of Helsinki. All tissue utilized for this study was acquired in the laboratory within 8 hours of death and processed immediately, and samples used for transcriptomic studies were processed within 5 hours of death. Human donor tissue was utilized in one scRNA-seq experiment using four eyes from four donors, and additional immunohistochemistry (IHC) experiments using eleven eyes from eleven donors. All eyes were originally selected as control eyes with no known history of ocular disease based on chart review, however donor “1” was subsequently found to have histopathological evidence of AMD.

### Single-cell RNA sequencing

#### Sample preparation

Macular RPE/choroidal tissue was isolated from donors using 8- or 12-mm punch biopsies centered on the fovea. Retinal tissue was removed, and remaining RPE/choroid was minced into 1 mm × 1 mm squares. Cells were placed in 2 mg/mL of collagenase II (Gibco) and resuspended in Hanks’ Balanced Salt Solution with calcium chloride and magnesium chloride (Life Technologies). Samples were placed on a 37° C shaker for 1 h to dissociate, then centrifuged and resuspended in Recovery Cell Culture Freezing Media (Life Technologies). Samples were then cooled at 1° C/min for 3-12 h in a -80° C freezer using a CoolCell LX container (Corning), and transferred to liquid nitrogen for long term storage.

#### Antibody labeling

Cryopreserved cells were quickly thawed in a 37° C water bath and labeled with barcoded antibodies according to the 10X Genomics Cell Surface Protein Labeling protocol outlined in document CG000149. 3.5 mL of PBS (Gibco) + 1 % Ultrapure BSA (Thermo Fisher Scientific) was added to each sample, and cells were spun at 4° C for 7 min @ 300 rcf. Supernatant was removed, and the cells were resuspended in 50 µL of cold PBS + 1% BSA, and incubated after addition of 5 µL Fc receptor blocker (Human TruStain FcX, BioLegend) for 10 minutes. An antibody cocktail was prepared by combining enough of each antibody to reach 1 µg of each per sample, centrifuged at 4° C for 10 min @ 14,000 rcf, then the antibody cocktail supernatant was transferred to a new tube. The antibody cocktail was subsequently added to the cells, then more cold PBS + 1 % BSA was added up to a total volume of 100 µL, and cells were incubated at 4° C for 30 minutes in the dark. Cells were then washed with 3.5 mL cold PBS + 1 % BSA and spun at 4° C for 7 min @ 300 rcf a total of 3 times. Cells were finally resuspended in 100 µL PBS + 1 % BSA, and counted with a hemocytometer. Cell suspensions were then lysed, barcoded, and libraries were generated according to the Chromium GEM-X Single Cell 3’ protocol with Feature Barcode technology for Cell Surface Protein (10X Genomics). Gene expression and surface protein libraries were pooled at a 4:1 ratio. Libraries were subsequently sequenced on the NovaSeq6000 at the University of Iowa’s Genome Sequencing Core (Illumina).

Antibodies used in the scRNA-seq study included: anti-CD34, anti-CD45, anti-CD55, anti-CD59, anti-C5b-9, and isotype controls from BioLegend (Supplemental Table 1). For quantifying cell surface MAC labeling, barcoded antibodies were not commercially available, and a custom labeling was performed by BioLegend. Unlabeled monoclonal anti-C5b-9 antibodies (clone aE11) were obtained from Novus biologicals Cat # NBP1-05120 and 1 mg was conjugated to a sequence containing a 15 base pair barcode. A subset of these antibodies were simultaneously conjugated to AF488 (BioLegend). In order to assess whether the antibody still retained its activity following conjugation, immunohistochemistry of donor choroid was performed as described previously to assure that the labeling was unchanged [10].

#### Computational analysis of scRNA-seq data

Bcl2fastq software (Illumina) was used to generate FASTQ files from base calls, which were mapped to human reference genome hg38 with CellRanger using the University of Iowa’s Argon High Power Computing (HPC) system. Cells with fewer than 2,000 or more than 7,000 genes expressed were filtered out, as well as cells with greater than 10% mitochondrial transcripts. Filtered libraries were normalized (gene expression using LogNormalize, antibody tags using CLR) and integrated with canonical correlation analysis (CCA) using the Seurat package. Fully processed scRNA-seq data are hosted online at https://singlecell-eye.org under the dataset “Transcriptomic Characterization of Terminal Complement Complex-bound Cells in Human Choroid Using Single-Cell RNA Sequencing” [18].

#### Immunohistochemistry

Flash-frozen, paraformaldehyde fixed sections of human retina and choroid were used for immunohistochemistry (IHC). Sections were blocked with 1 mg/mL BSA for 15 minutes before incubation with primary antibodies for one hour. Sections were subsequently incubated with secondary antibodies resuspended in PBS with diamidino-2-phenylindole (DAPI) for 30 minutes (Sigma). These antibodies are listed in Supplemental Table 2. Whole mounts were prepared by removing RPE/choroid from the sclera, then carefully scraping away overlying RPE. Choroidal tissue was blocked, and incubated in primary antibodies for four hours, and in secondary overnight. These antibodies are listed in Supplemental Table 3. Both sample types were washed three times in PBS, mounted using Aqua-Mount Mounting Medium (Thermo Fisher Scientific) and then visualized using a Leica TCS SPE upright confocal microscope system (Leica Microsystems).

## 2. Results

Samples from four human donor eyes were used for the scRNA-seq experiment. Samples were chosen from aged donors without clinical history of AMD, although one donor had histopathological evidence of AMD noted on postmortem H&E. Donor information is provided in Table 1. An overview of the recovered choroidal cell types is shown in Figure 1A, and represents the expected cell types based on prior scRNA-seq studies in the human choroid [19]. Choroidal cell types were annotated according to gene expression profiles in the literature, and Figure 1B illustrates several genes used to annotate the different clusters. Cells were binned into 17 clusters ranging from 23 to 1891 cells. Full names of cell clusters are listed in Table 2. 8644 cells were recovered across all four donor samples in this experiment, with appreciable contributions from each donor (Figure 1C).

**Table 1.** Information for the four human donors included in the scRNA-seq experiment.

| Donor | Age | Sex | Cause of death | Time to preservation | Ophthalmic notes |
| --- | --- | --- | --- | --- | --- |
| 1 | 84 | F | Cardiac arrest | <5 hours | No ophthalmic records, histopathological evidence of AMD |
| 2 | 76 | F | Cardiogenic shock | <5 hours | No ophthalmic records, no gross pathology |
| 3 | 77 | M | Subarachnoid hemorrhage | <5 hours | No ophthalmic records, unremarkable for drusen on histology |
| 4 | 83 | F | Pneumonia | <5 hours | History of diabetes without retinopathy, unremarkable for drusen on histology |

**Table 2.** Full names corresponding to the abbreviated cell type names in the scRNA-seq experiment.

| Cell Type Shorthand | Full Name |
| --- | --- |
| RPE | Retinal pigment epithelial cell |
| Schwann | Schwann cell |
| Artery_Endothelial | Arterial endothelial cell |
| Vein_1_Endothelial | Venous endothelial cell, cluster 1 |
| Vein_2_Endothelial | Venous endothelial cell, cluster 2 |
| BCell | B cell/B lymphocyte |
| CD8_T | CD8+ T lymphocyte |
| CD4_T | CD4+ T lymphocyte/T helper cell |
| NK | Natural killer cell |
| Macrophage_Inf_1 | Infiltrating macrophage, cluster 1 |
| Macrophage_Inf_2 | Infiltrating macrophage, cluster 2 |
| Macrophage_Resident | Resident macrophage |
| Fibroblast | Fibroblast |
| Melanocyte | Melanocyte |
| Pericyte | Pericyte |
| SMC | Vascular smooth muscle cell |

**Figure 1.**
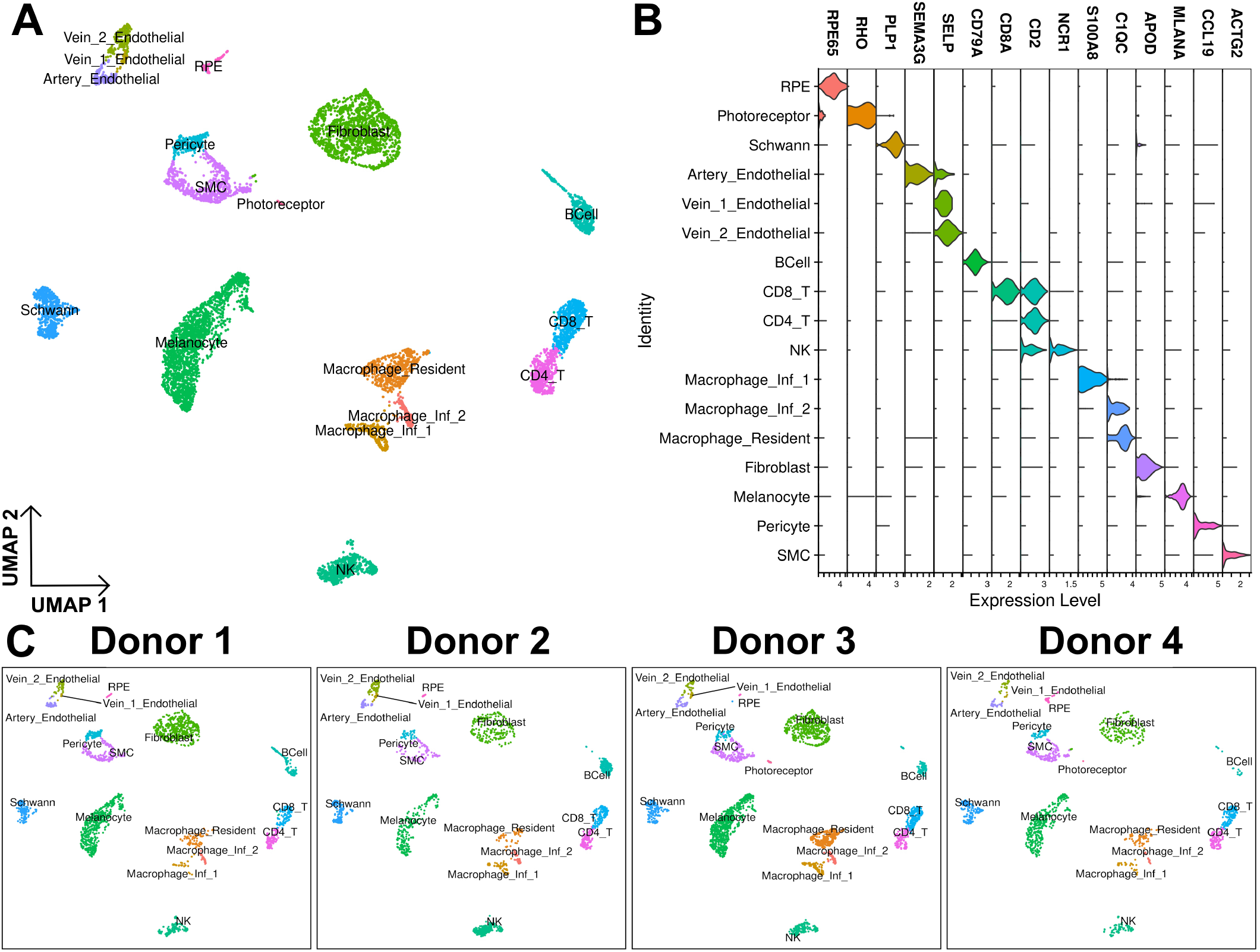
Sample overview. An overview of cell types recovered in this scRNA-seq experiment is shown in (A). A summary of genes used to annotate clusters is shown in (B). Cells recovered across each donor is shown in (C).

In order to detect and quantify cell surface proteins at single cell resolution, we labeled our cells with antibodies directed against surface proteins of interest conjugated with single-stranded DNA barcodes. Using a cell surface protein-compatible scRNA-seq protocol, the antibodies are able to be detected on a per-cell basis in the sequencing data. To evaluate the distribution of complement and its regulators in the choroid, we included antibodies directed against complement inhibitors CD55 and CD59, as well as an antibody directed against the assembled MAC. We also included antibodies directed against CD34 and CD45 to detect endothelial and white blood cells, respectively. Finally, we included barcoded isotype control antibodies (e.g. mouse IgG1, mouse IgG2a) which share a constant region with our experimental antibodies to establish the level of background signal across each cell type. Plots showing functionality of the barcoded CD45 antibody are shown in Figure 2, while plots for CD34, CD55, and CD59 antibodies are shown in Supplemental Figure 1. The MAC antibody, which has no corresponding gene expression data, is discussed below.

**Figure 2.**
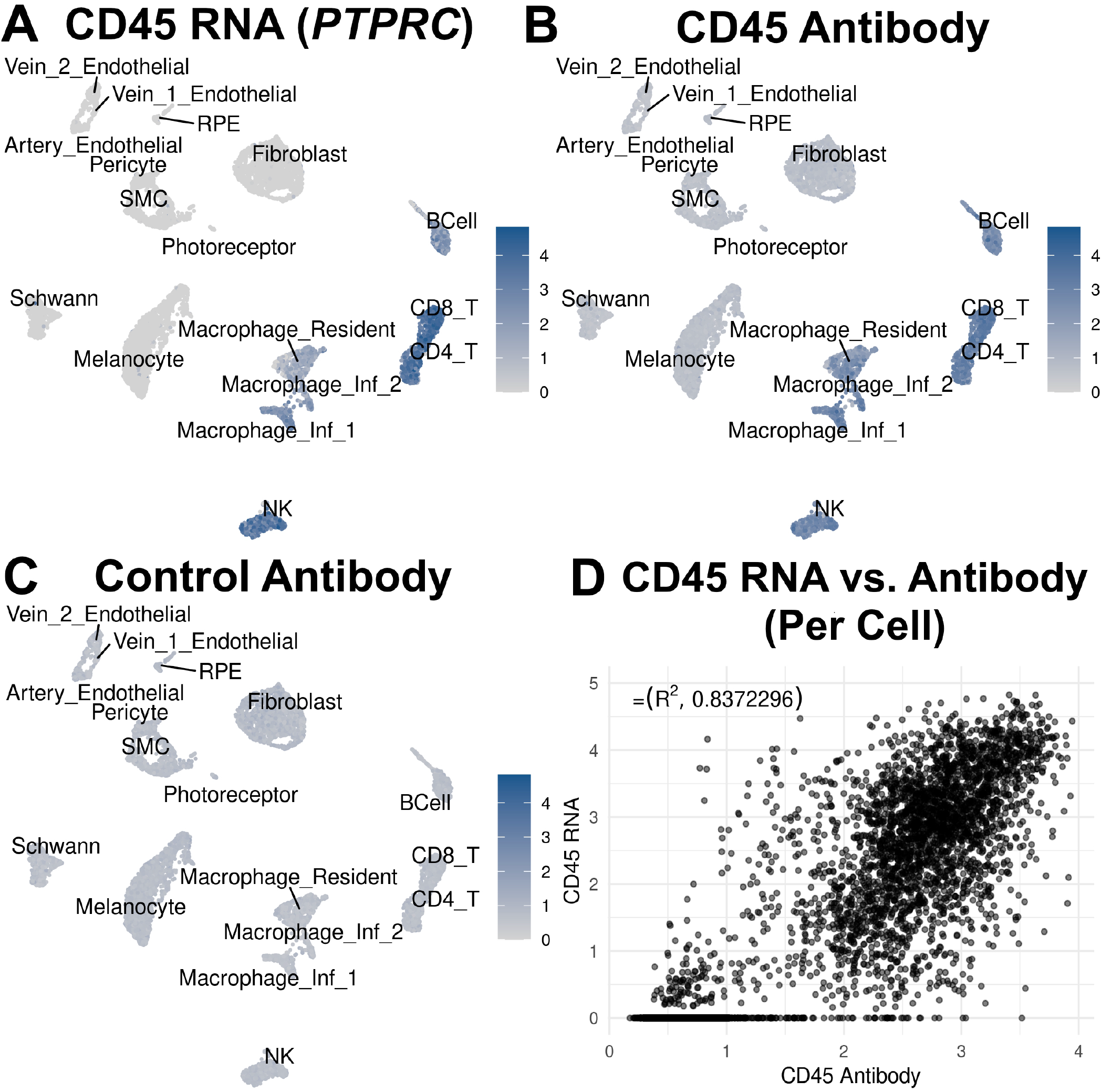
Antibody overview. A plot showing expression of CD45 (*PTPRC*) RNA across recovered cell types is shown in (A). Plot showing CD45 antibody detection across recovered cells is shown in (B), while the same plot showing isotype control antibody (Mouse IgG1) detection is shown in (C). (D) shows a scatterplot of antibody detection vs. corresponding RNA across all cells on a per-cell basis.

To evaluate the specificity of the antibody detection, we assessed the pattern of detection across our recovered cell types. We expected CD45 gene expression to be highest in leukocyte clusters, with CD45 surface protein detection mirroring this trend. We observed elevated CD45 (*PTPRC*) gene expression exclusive to leukocyte clusters (Figure 2A), as well as strong CD45 antibody signal above background that was similarly exclusive to those leukocyte clusters (Figure 2B, 2C). We saw a similar pattern of congruent *CD34* gene expression and antibody signal exclusive to endothelial cells (Supplemental Figure 1A, 1B, 1C).

We also quantified binding and gene expression patterns of complement inhibitors CD55 and CD59 by cell type, expecting the signal to be most abundant in non-circulating cells closely exposed to circulating complement, endothelial cells and pericytes. Ultimately, we found *CD55* gene expression in a variety of cell types, especially in leukocytes, but CD55 surface protein binding was comparatively limited, and greatest in melanocyte and Schwann cell clusters (Supplemental Figure 1E, 1F, 1G). *CD59* gene expression and surface protein were detected in endothelial cells and pericytes as expected, but also frequently detected on melanocytes, fibroblasts, and Schwann cells (Supplemental Figure 1I, 1J, 1K).

In addition to specificity, we evaluated the relationship between normalized “expression level” of the antibody signal and the level of its corresponding mRNA. The protein and RNA levels varied widely for different gene products. For example, we saw a strong correlation between our normalized CD45 (*PTPRC*) gene expression vs. normalized CD45 surface protein on a per-cell basis as shown in Figure 2D. We show weaker, albeit slightly positive correlations for CD34 and CD59 antibody vs. RNA (Supplemental Figures 1D and 1L), while CD55 antibody binding and RNA are very weakly correlated (Supplemental Figure 1H), suggesting the fluctuations in the relationship between gene expression and surface detection among these different protein species reflects regulation at different levels.

The MAC has been shown to deposit into cell membranes in quantities that can trigger intracellular effects downstream without causing lysis of the cell [20]. To that end, we attempted to identify the MAC on the surface of our recovered choroidal cells using a barcoded antibody against the assembled complex. In order to account for cells with high background signal (e.g. RPE, fibroblasts) identified using the isotype control antibody (Mouse IgG2a), we set a conservative threshold for MAC surface “positivity” at the maximum value of the corresponding control antibody in each individual cell type. Normalized but otherwise unmodified detection values across all cell types are shown in Supplemental Figure 2. We then binned cells with MAC detection above this threshold as “positive”, and arranged cell types by decreasing fraction in Figure 3A. We saw few “MAC positive” cells across most recovered choroidal cell types, with a few exceptions: vascular mural cells and members of the macrophage family. Based on prior immunohistochemical studies in the human choroid, we anticipated that MAC would be present primarily on the cells of the choriocapillaris, which has been robustly shown to label positively with MAC in aged donors [10]. Unfortunately, this experiment did not capture significant quantities of capillary endothelial cells. Additionally, the RPE (a cell type of interest in the context of early AMD), was captured in a moderate quantity but showed very little surface MAC above background. However, nearly 10% of recovered pericytes, supportive vascular mural cells of the choriocapillaris, labeled positively for MAC in the single cell experiment. A similar fraction of vascular smooth muscle cells originating from the deeper vessels labeled positively as well. Immunohistochemistry experiments on choroid from 11 aged donors showed modest MAC colocalization with pericytes in the choriocapillaris, with a representative en face immunolabeled image in Figure 3B. While we did calculate Spearman correlations for expressed genes vs. the MAC in these mural cells, we were not able to detect any genes that appeared to scale convincingly with surface MAC in this experiment.

**Figure 3.**
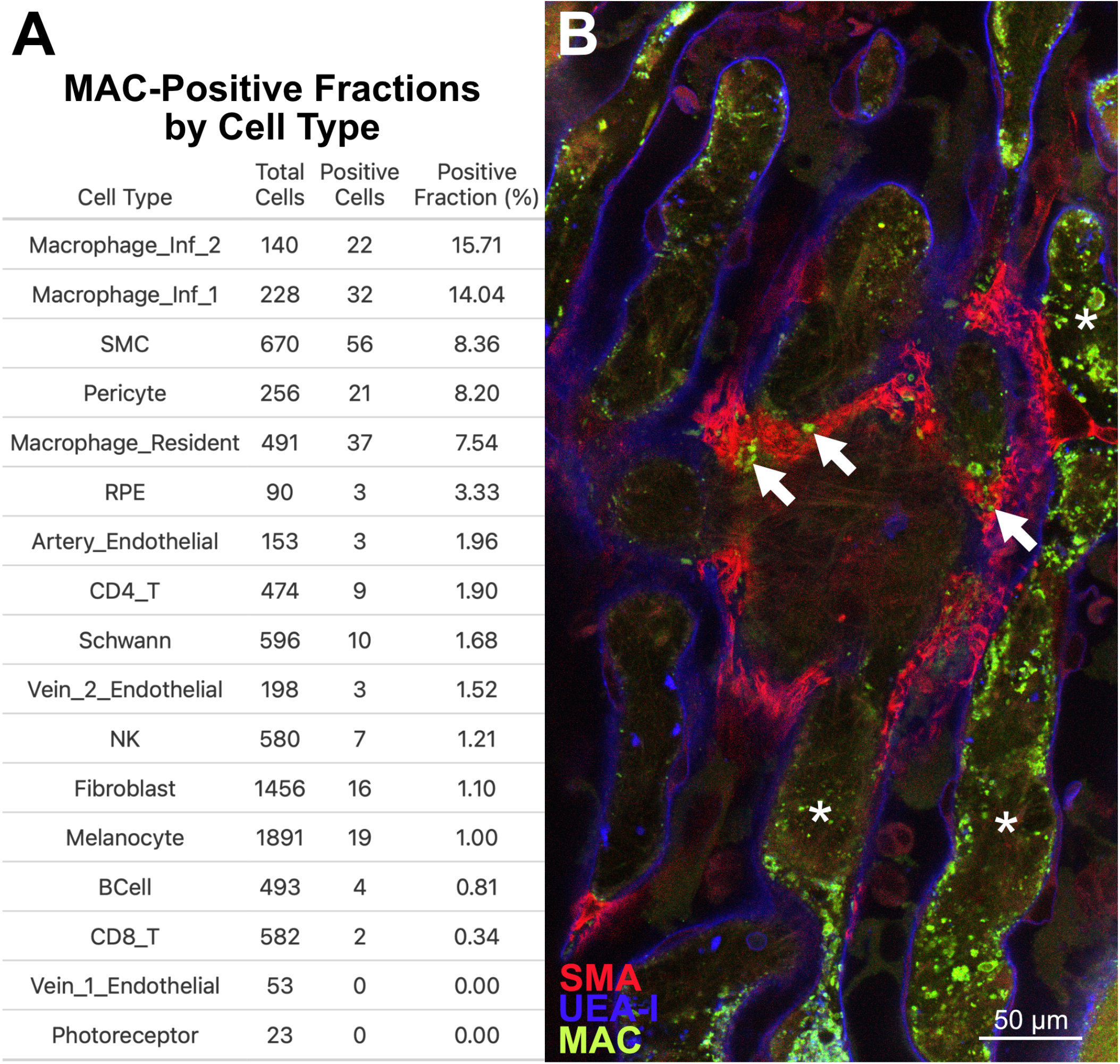
MAC detection. Membrane attack complex (MAC) detection across recovered cell types is shown in (A) in descending order by positive fraction. An image of human choriocapillaris labeled for smooth muscle (SMA, red), endothelial cells (UEA-I, blue), and the assembled MAC (green). While MAC is predominantly associated with endothelial cells and intercapillary pillars (asterisks), MAC punctae are also associated with some pericytes (arrows) (B).

Although our initial goal was to assess vascular cells, members of the macrophage family had the highest proportion of cells with surface MAC above background. The two populations of infiltrating macrophages had the greatest MAC-positive fraction in this experiment (∼15%). Resident macrophages also had an appreciable positive fraction (∼8%), with a sharp dropoff thereafter. We did note that surface MAC appeared to be higher in the donor with the least surface CD59 on macrophages (Supplemental Figure 3A, 3B). The MAC has been shown to activate membrane phospholipases to trigger downstream changes in gene expression and protein synthesis, eventually resulting in increased proliferation and protection from apoptosis [20, 21]. Macrophages are also thought to be activated by sublytic MAC, with differing roles in different tissues. Macrophages have demonstrated NLRP3 inflammasome activation and increased IL-18 in response to sublytic MAC [22]. In dendritic cells, sublytic MAC may promote dendritic cell maturation, in turn promoting T cell Th1 polarization [23].

We then turned our focus to the 404 endothelial cells recovered across the four donors. We recovered arterial and venous endothelial cells from each donor (Figure 4A). Although our analysis revealed three endothelial cell clusters, none of the clusters convincingly represented endothelial cells in the choriocapillaris. These cells, which show the highest degree of MAC colocalization in immunohistochemical studies, also show high expression of *CA4* at both protein and RNA levels [19, 24]. However, a *CA4*-positive cluster was not identified among our recovered cells. Although there was little MAC on the surface of our recovered arterial and venous endothelial cells (Figure 3A), complement regulators CD55 and CD59 were detected on the surface of all endothelial cell subtypes.

**Figure 4.**
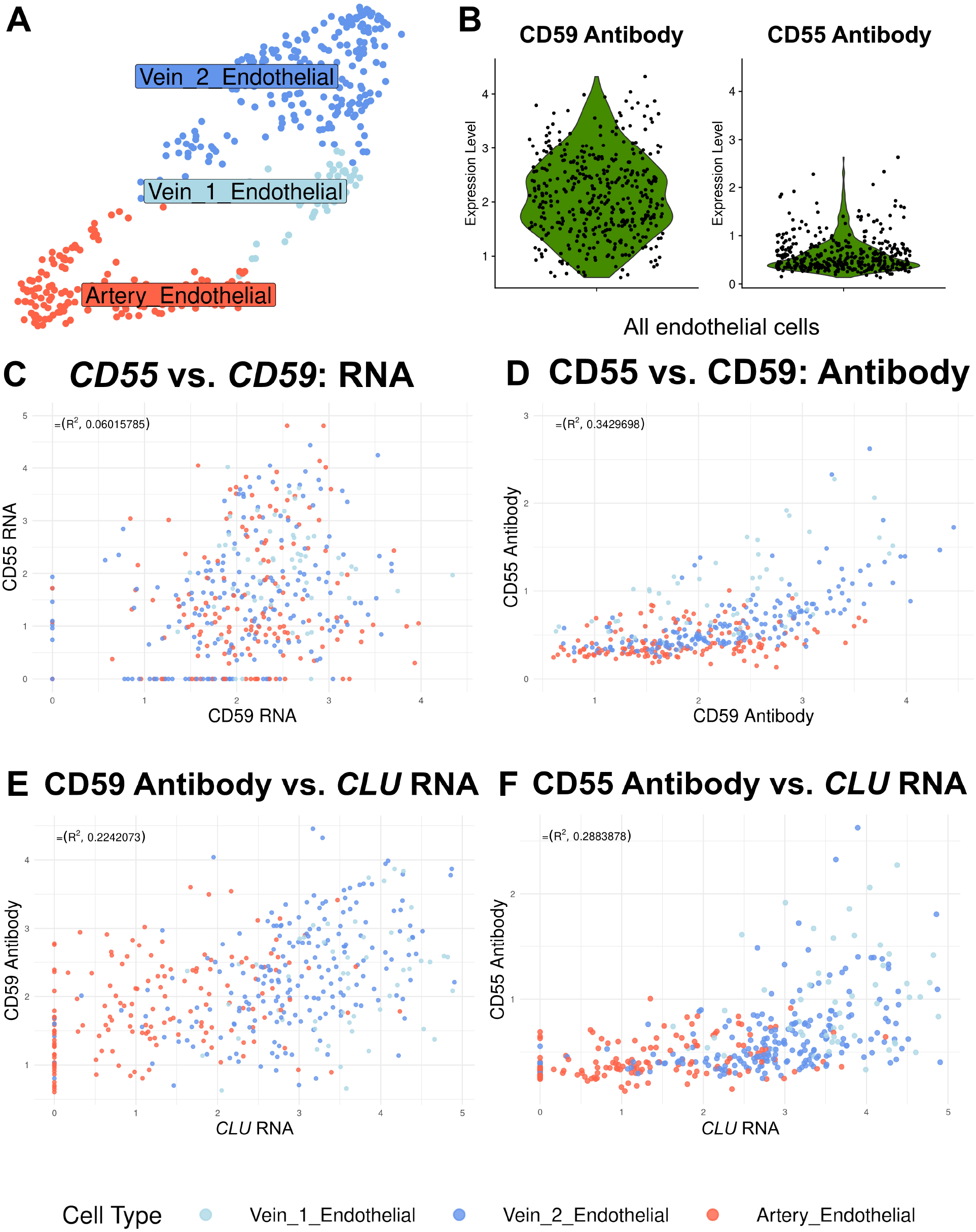
Endothelial cells & complement regulators. (A) Overview of recovered endothelial cells. (B) Violin plots illustrating continuous CD55 and CD59 surface detection. (C) Scatterplot showing *CD55* and *CD59* gene expression on a per cell basis, with cells colored by cell type. (D) CD55 and CD59 cell surface detection. CD55 (E) and CD59 (F) vs. *CLU* gene expression.

Interestingly, we saw no relationship between CD55 and CD59 at the gene expression level (Figure 4C). However, there was a moderate positive correlation between CD55 and CD59 surface protein, suggesting that cells with a greater abundance of one surface-bound complement regulator have more of the other (Figure 4D). Furthermore, those cells with more surface CD55 and CD59 tended to be venous. This positive correlation was not present while comparing CD55 or CD59 surface binding with isotype controls (Supplemental Figure 4A, 4B). In order to assess the degree to which other complement regulators might be associated with surface CD55 or CD59, we calculated Spearman correlations across gene expression with abundance of these surface proteins. Among the genes most correlated with surface CD55 or CD59 was *CLU*, encoding clusterin, a secreted complement inhibitor that binds to assembling MAC preventing it from becoming active (4E, 4F) [25]. Interestingly, another gene with expression weakly correlated with CD55 and CD59 surface protein was *SGMS2* (Supplemental Figure 5A, 5B). *SGMS2* encodes sphingomyelin synthetase 2, an enzyme residing at the plasma membrane which synthesizes the components of the lipid rafts that hold GPI-anchored proteins at the surface, like CD55 and CD59 [26, 27].

## 3. Discussion

In the past, we have used scRNA-seq to characterize the cell types and gene expression patterns in the human choroid [16, 19]. In the present study, we were able to expand on these experiments by showcasing a multiomic approach to combine gene expression analysis with cell surface protein detection. We were able to detect the membrane attack complex (MAC), as well as complement regulators CD55 and CD59, and other cell surface proteins (CD34 and CD45) on the surface of cells isolated from human choroid.

One of our goals was to measure surface MAC deposition across the different cell types in the choroid using an unbiased approach. While we were able to do so, we originally anticipated that the MAC would be limited to the endothelial cells of the choriocapillaris. Interestingly, the cell types with the greatest proportion of surface MAC in this experiment were the members of the macrophage family, pericytes, and smooth muscle cells. We were able to follow these results up by detecting MAC on the surface of pericytes and smooth muscle cells across numerous donors by imaging, suggesting that the MAC may be depositing on more cell types than expected. Still, the impact of MAC deposition on these cells, both short and long-term, remains unclear. Ultimately, these findings point to the MAC playing multiple roles in the choroid, accumulating in sublytic quantities on various different cell types.

This study does have several limitations. While we recovered several hundred endothelial cells and pericytes in our single cell experiment, we recovered almost no *CA4*-expressing endothelial cells, far too few to form a convincing capillary endothelial cell cluster. The ratio of capillary endothelial cells to pericytes in tissue is approximately 6:1 based on the histological literature, whereas in our study we captured little to no capillary endothelial cells and 256 pericytes, suggesting that our recovered cells are depleted for capillary endothelial cells [28].

The depletion of choriocapillaris endothelial cells may contribute to the decreased MAC signal seen in our single cell experiment as compared to human tissue imaged after immunohistochemistry, in addition to MAC deposition on extracellular matrix. It is possible that capillary endothelial cells are more difficult to isolate from the matrix, or are less tolerant to the extensive tissue digestion, cryopreservation, thawing, and antibody labeling steps prior to capture. Either way, past single cell experiments from our group also found that capillary endothelial cells were underrepresented in recovered choroidal cells, and it appears that enrichment or in-vitro modeling will likely be required for future experiments of this type focused on the choroidal endothelium [19].

From the endothelial cells that were captured, we found that arterial and venous subtypes featured CD55 and CD59 detectable at the cell surface and by mRNA expression across all donors, and that the arterial endothelial cell population featured less than the venous populations. Additionally, *CLU* expression was correlated with surface CD55 and CD59, suggesting that venous endothelial cells feature greater complement protection than arterial cells by several different mechanisms. Notably, *CLU* expression was also highest in the venous endothelial cells in Voigt et al. 2022 [19]. Ultimately, these data suggest that complement protection is heterogeneous across choroidal endothelial cell subtypes, with venous endothelial cells featuring a greater abundance of complement regulators detectable at the RNA and protein levels.

Pericytes showed an unexpected level of anti-MAC labeling. While these cells have not been the primary focus of research in early AMD and dysfunction in the aging choroid, there is reason to believe they may play a role in the pathophysiology of other ocular and extraocular vascular diseases. Interestingly, pericyte susceptibility to complement-mediated damage has been considered as a mechanism contributing to pericyte loss in the retinal vasculature in diabetic retinopathy, although the inciting pathophysiology and histopathological changes in early AMD are considerably different [29, 30]. Terminal complement may also damage smooth muscle-like cells in the kidney microvasculature in the context of several forms of glomerulonephritis [31]. These interesting findings suggest further exploration of pericytes in the context of AMD pathogenesis.

Overall, this dataset provides novel insight into the patterns of MAC deposition and select complement regulators across the various cell types in the aging human choroid.

## 4. Other/Acknowledgements

- Funded in part by NIH Grants EY-024605, EY-025580, and T32GM145441.
- The authors thank the Iowa Lions Eye Bank for their partnership in tissue procurement and gratefully acknowledge the eye donors and their families for their generous and invaluable contribution to biomedical research.

