## Supplemental Figures for "Transcriptomic Characterization of Terminal Complement Complex-bound Cells in Human Choroid Using Single-Cell RNA Sequencing"

### 1. Supplemental Figures

**Supplemental Table 1:**
**Single Cell RNA Sequencing Antibodies:**

| BioLegend Part No. | Target | Isotype | Conjugation | Labeling Concentration | Barcode Sequence |
|---|---|---|---|---|---|
| 343539 | CD34 | Mouse IgG1 κ | TotalSeq-B0054 | 1 µg/100 µL | GCAGAAATCTCCCTT |
| 304066 | CD45 | Mouse IgG1 κ | TotalSeq-B0391 | 1 µg/100 µL | TGCAATTACCCGGAT |
| 311321 | CD55 | Mouse IgG1 κ | TotalSeq-B0383 | 1 µg/100 µL | GCTCATTACCCATTA |
| 304713 | CD59 | Mouse IgG2a κ | TotalSeq-B0361 | 1 µg/100 µL | AATTAGCCGTCGAGA |
| 900009786 | C5b-9 (MAC) | Mouse IgG2a κ | TotalSeq-B5101/AF488 | 1 µg/100 µL | TACTCACCGTAGTAC |
| 359139 | CD195 | Rat IgG2b κ | TotalSeq-B0141 | 1 µg/100 µL | CCAAAGTAAGAGCCA |
| 304066 | Isotype Control | Mouse IgG1 κ | TotalSeq-B0090 | 1 µg/100 µL | GCCGGACGACATTAA |
| 400185 | Isotype Control | Mouse IgG2a κ | TotalSeq-B0091 | 1 µg/100 µL | CTCCTACCTAAACTG |
| 400689 | Isotype Control | Rat IgG2b κ | TotalSeq-B0095 | 1 µg/100 µL | GATTCTTGACGACCT |

**Supplemental Table 2:**
**Labeling Molecules for Immunohistochemistry (Choroidal sections):**

| Manufacturer | Labeling Molecule | Molecule Type | Host Species | Labeling Concentration | Tag |
|---|---|---|---|---|---|
| Abcam | Anti-C5b-9 | Antibody | Rabbit | 16.67 µg/mL | N/A |
| Novus Biologicals | Anti-C5b-9 | Antibody | Mouse | 3.33 µg/mL | N/A |
| Millipore-Sigma | Anti- α-smooth muscle actin | Antibody | Mouse | 3.75 µg/mL | Cy3 |
| Thermo Fisher | Anti-rabbit | Antibody | Donkey | 10 µg/mL | AF488 |
| Thermo Fisher | Anti-mouse | Antibody | Donkey | 10 µg/mL | AF488 |
| Vector Laboratories | UEA-I | Lectin | *Ulex europaeus* | 20 µg/mL | Biotin |
| Vector Laboratories | Texas Red | Fluorophore | N/A | 10 µg/mL | Streptavidin |

**Supplemental Table 3:**
**Labeling Molecules for Immunohistochemistry (Choroidal whole mounts):**

| Manufacturer | Labeling Molecule | Molecule Type | Host Species | Labeling Concentration | Tag |
|---|---|---|---|---|---|
| Abcam | Anti-C5b-9 | Antibody | Rabbit | 16.667 µg/mL | N/A |
| Millipore-Sigma | Anti- α-smooth muscle actin | Antibody | Mouse | 3.75 µg/mL | Cy3 |
| Thermo Fisher | Anti-rabbit | Antibody | Donkey | 10 µg/mL | AF488 |
| Vector Laboratories | UEA-I | Lectin | *Ulex europaeus* | 20 µg/mL | Biotin |
| Vector Laboratories | DyLight 649 | Fluorophore | N/A | 10 µg/mL | Streptavidin |

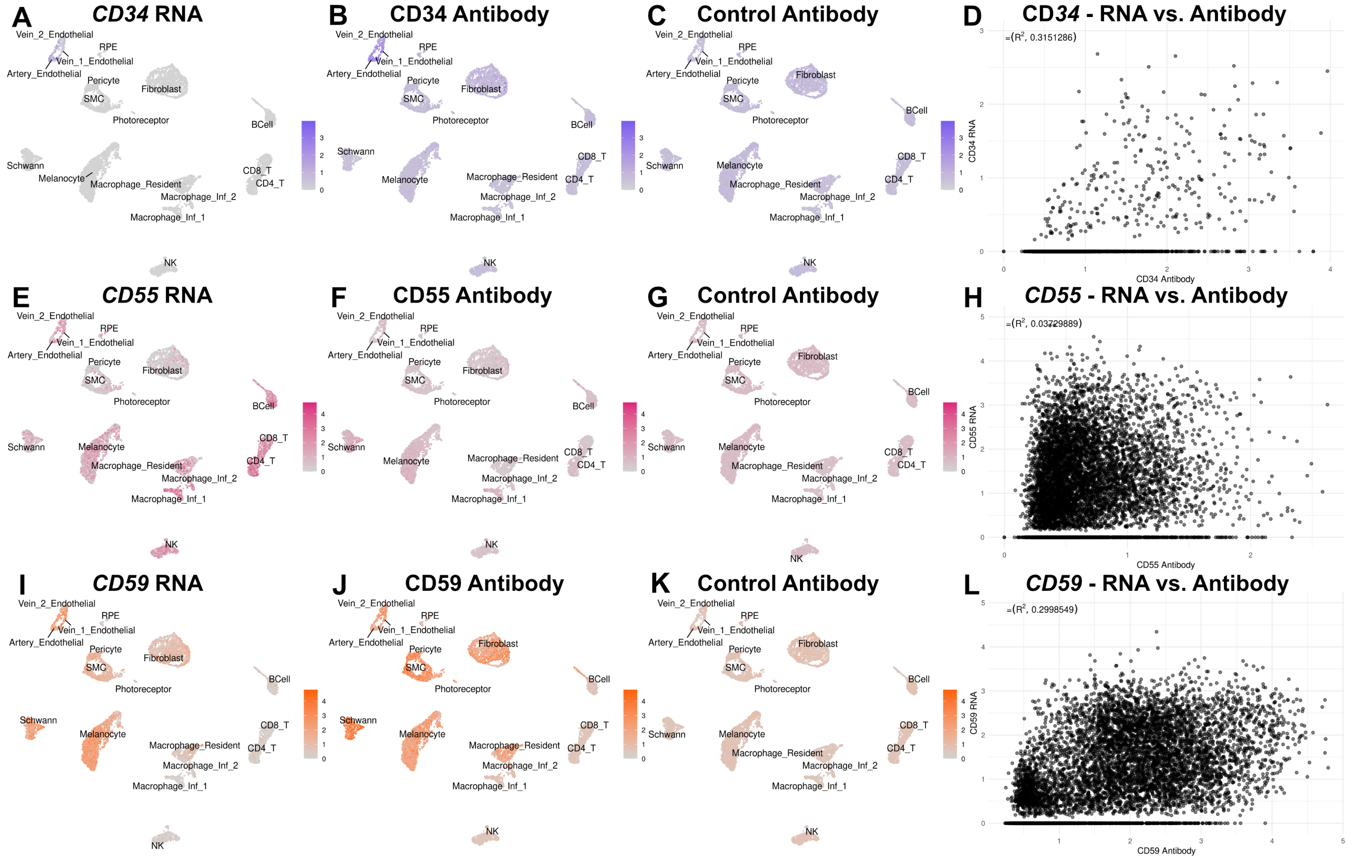


**Supplemental Figure 1. CD34, CD55, CD59 surface detection across all cells.** Gene expression plots for CD34, CD55, and CD59 are shown in (A), (E), and (I), respectively. Corresponding antibody signals are shown in (B), (F), and (J). Isotype control signal is shown in (C), (G), and (K). Gene expression vs. antibody signal per cell is plotted across all cells for *CD34*, *CD59*, and *CD55* in (D), (H), and (L), respectively.

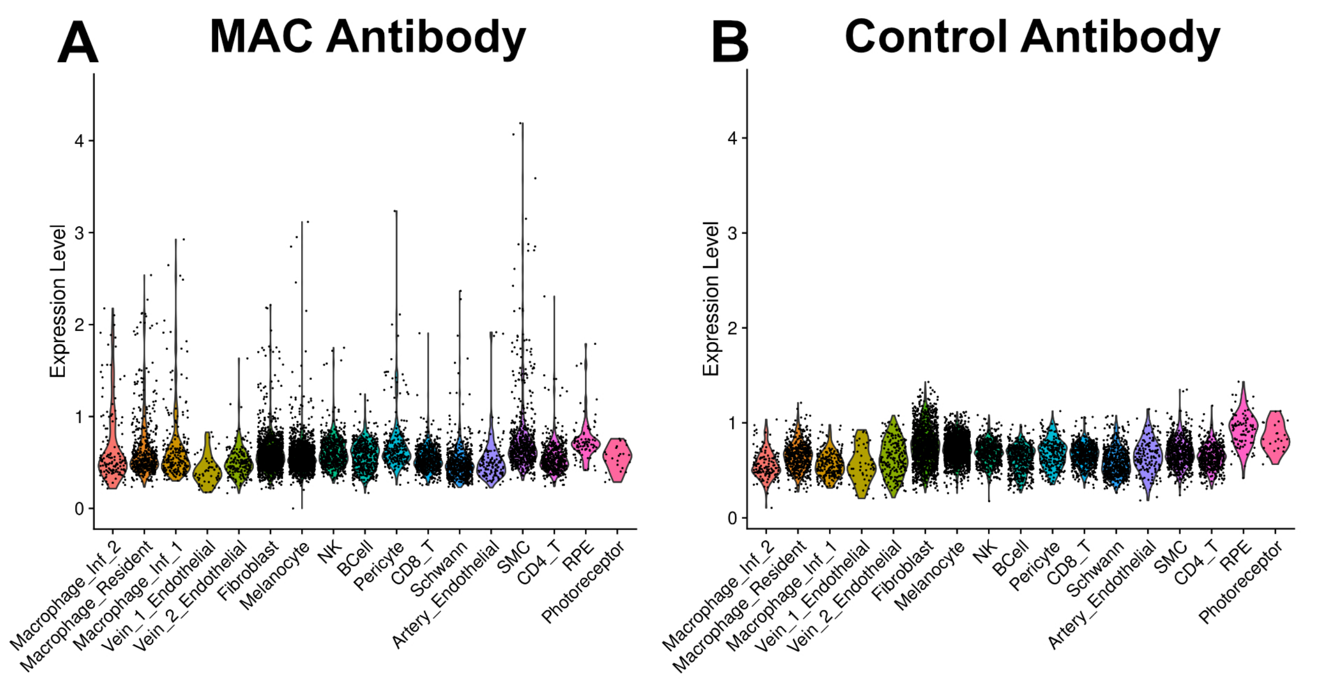


**Supplemental Figure 2. MAC detection across all cell types.** The MAC antibody signal across our recovered cell types is plotted in (A). The signal for the corresponding isotype control (IgG2a) in each cell type is shown in (B).

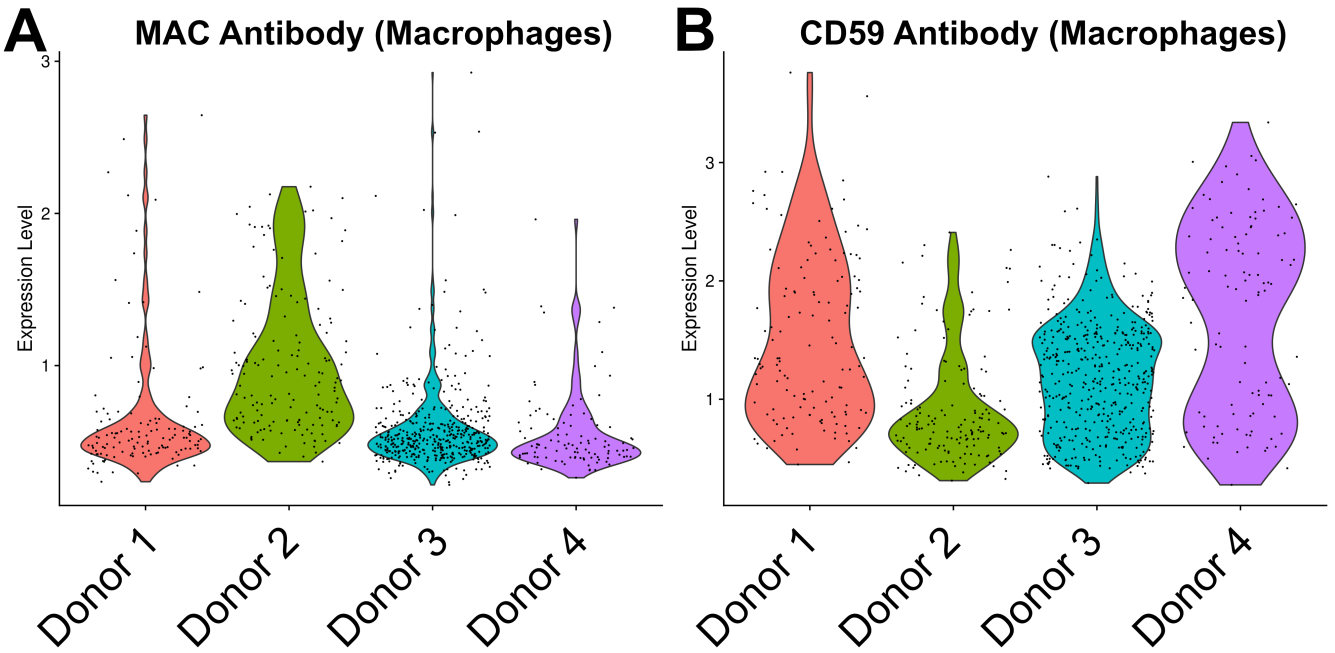


**Supplemental Figure 3. MAC on macrophages across donors.** MAC antibody detection on the macrophage family is plotted across all donors in (A). CD59 antibody detection is shown in (B) across the same donors.

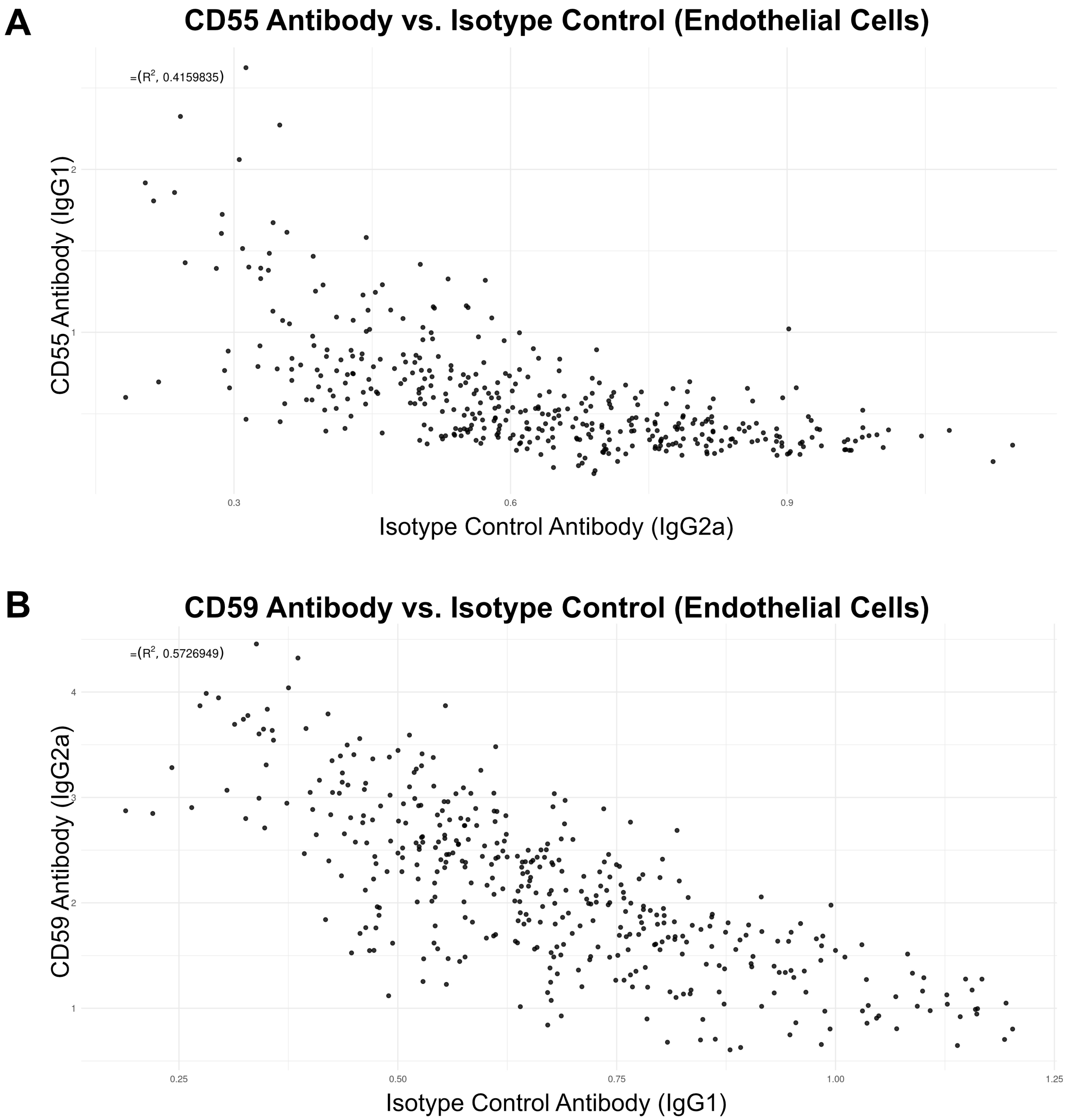


**Supplemental Figure 4. Complement regulators vs. isotype controls on endothelial cells.** A scatterplot of CD55 antibody (IgG1) detection vs. the CD59 antibody's isotype control (IgG2a) on a per-cell basis is shown in (A). A similar plot of CD59 antibody (IgG2a) detection vs. the CD55 antibody's isotype control (IgG1) is shown in (B).

**A**

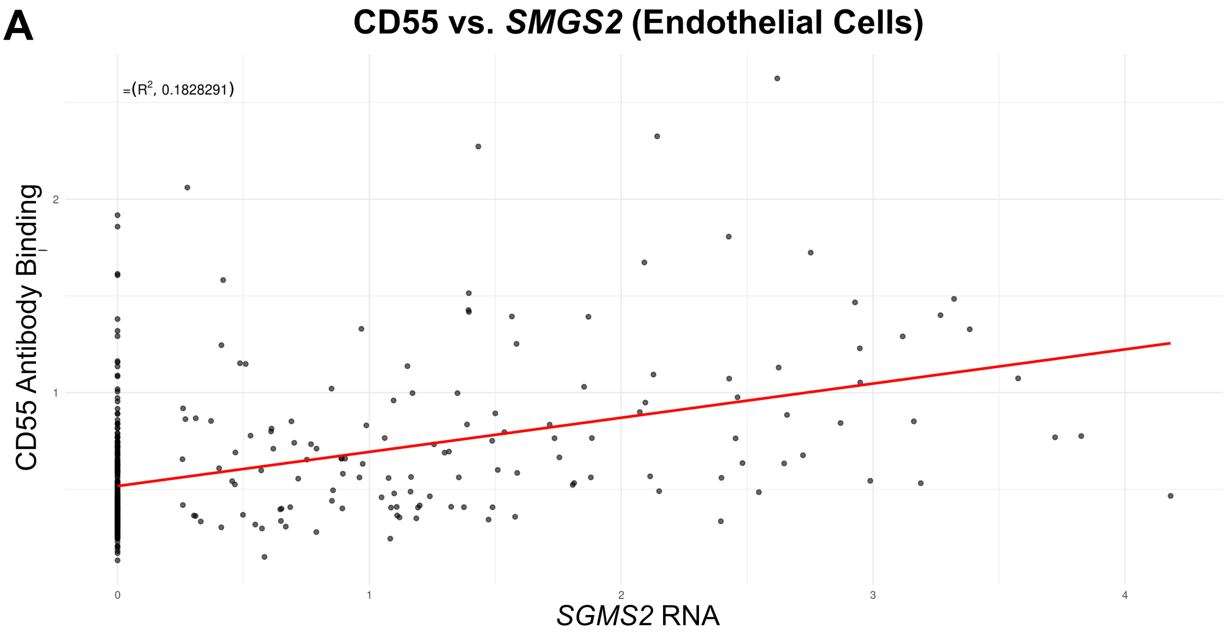


**B**

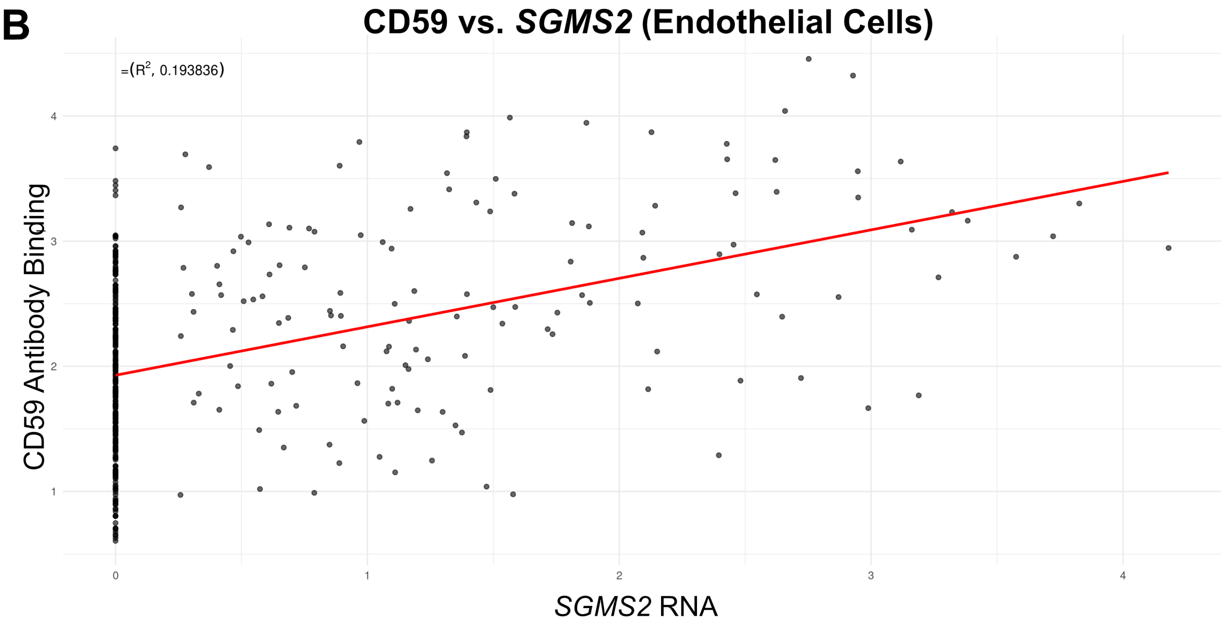


**Supplemental Figure 5. Complement regulator detection vs. *SGMS2* expression on endothelial cells.** A scatterplot showing CD55 antibody detection vs. *SGMS2* gene expression on a per-cell basis is shown in (A), while the same plot for CD59 is shown in (B).
